# Neuronal microscale biophysical instability mediates macroscale network dynamics shaping pathological manifestations

**DOI:** 10.64898/2026.01.20.697254

**Authors:** Vipin Kumar, Victor M. Sanchez Franco, Faith S. Ferry, Yizhou Xie, Anelise N. Hutson, Yutian J. Zhang, Skylar D. Daniels, Dieu Linh Nguyen, Lucia K. Spera, Eleanora M. Snyder, Anneke Knauss, Srilakshiya L. Sudhakar, Grace Y. Duan, Elizabeth M. Paul, Masashi Tabuchi

## Abstract

Subtle changes in membrane excitability may contribute to neurological disease, but disease-relevant dynamical signatures that generalize across models remain poorly defined. Here, we quantified variability in action potential initiation in *Drosophila* neurons expressing tauopathy- or epilepsy-associated mutations and in human iPSC-derived neurons from patients with Alzheimer’s disease or epilepsy. Across these models, disease-associated neurons exhibited increased instability in spike timing relative to controls. In *Drosophila* neurons, this phenotype was accompanied by increased variability in voltage-gated sodium currents during non-stationary inactivation, identifying a candidate biophysical contributor to altered spike initiation.

Antiepileptic drugs reduced sodium-current variability and stabilized spike initiation in fly neurons, and similarly improved spike-timing instability in patient-derived human neurons. In the fly models, neuronal instability was also associated with altered circuit- and brain-state readouts. Together, these findings identify unstable spike initiation as a conserved electrophysiological phenotype across distinct neurological disease models and suggest that sodium-channel-dependent variability may contribute to this phenotype. Rather than establishing a complete multiscale causal framework, our study defines a tractable cellular and dynamical entry point for investigating how subtle perturbations in intrinsic excitability may scale toward circuit dysfunction and disease-relevant phenotypes.

**Significance Statement:** Linking microscale neuronal changes to macroscale disease phenotypes remains a key challenge in neuroscience biophysics. Here, we show that neurons from *Drosophila* models of tauopathy and epilepsy and human iPSC-derived neurons from patients with Alzheimer’s disease and epilepsy share increased biophysical instability in their local neural activities. In fly neurons, this phenotype is associated with increased variability in voltage-gated sodium currents and is reduced by antiepileptic treatment. These findings define unstable local spike variability as a conserved dynamical signature across distinct disease models and nominate sodium-current variability as a mechanistically testable, pharmacologically reversible contributor to pathological excitability.

## Introduction

Our understanding of microscale and macroscale dynamics in biological systems remains limited, making it challenging to establish mechanistic links between microscale interactions and emergent macroscale behaviors, particularly when nonlinear, stochastic, and multiscale processes interact across levels. Understanding this connection is not only fundamentally important but also clinically significant, as it reveals how changes at the microscale translate into pathological outcomes at the macroscale, providing insights into disease progression. For example, identifying molecular targets critical to disease development by dissecting microscale and macroscale dynamics can aid in the development of early diagnostic biomarkers and targeted therapeutic strategies. Despite the recognition of the importance of this goal, bridging the gap between biological system hierarchies has been rare. By exploring the microscale dynamics within neurons, we gain profound insights into the subtle but powerful “pathological” biophysical changes that underlie macroscale pathological manifestations. This exploration is a critical step in unraveling the complexities of disease progression by shedding light on the nuanced changes that occur at the molecular and cellular levels. For this exploration, we examine the biophysical origins of the altered neuronal dynamics observed in tauopathy and epilepsy. To unravel the interplay between microscale changes and the resulting macroscale consequences, we utilize a specific circadian neuron expressing human mutant tau (Tau4RΔK) to model tauopathy, and a paralytic voltage-gated sodium channel gene mutation to model epilepsy (*para*^*bss*^, a bang-sensitive mutant) *(Parker et al., 2011)*. Voltage-gated sodium channels are crucial in shaping neuronal membrane potential dynamics (Catterall et al., 2005), especially during the initial depolarization phase of action potential generation, which affects neuronal firing patterns (Marban et al., 1998; Rutecki, 1992). Thus, we hypothesize that the expression of tauopathy and epilepsy-associated molecules would disrupt sodium channel function, leading to increased variability in action potential initiation. We show that the dynamical instability of the spike onset rapidness in DN1p circadian clock neurons in *Drosophila* was caused by Tau4RΔK and *para*^*bss*^ to increase local irregularity of DN1p spiking patterns through increased noise variability of voltage-gated sodium channels during their non-stationary inactivation process. We find that administering antiepileptic medications to suppress neuronal dynamical instability stabilizes not only the microscale biophysical properties of membrane potential in individual neurons but also the macroscale organization of global brain activity. Finally, we find that human iPSC-derived neural cultures from Alzheimer’s disease (AD) patients show similar alteration of neuronal dynamics, which are also sensitive to the presence of antiepileptic drugs. Our study bridges the gap between microscale biophysical dynamics and macroscale system dynamics in the context of pathological molecule perturbations, revealing a shared mechanism by which AD and epilepsy-related molecules disrupt reliable spike trains and cause pathological modifications, ultimately leading to disease-associated behavioral outcomes.

## Results

We employed a DN1p circadian network in *Drosophila* expressing the human mutant tau (Tau4RΔK), a well-established tauopathy-associated factor due to its pathological tau aggregation, and the *para*^*bss*^, an allele of the paralytic (para) voltage-gated sodium channel gene that exhibits seizure-like behaviors and serves as a model for epilepsy. These systems allowed us to investigate how microscale perturbations within a small population of neural clusters translate into macroscale outcomes in global brain state having neurodegenerative and seizure-related pathologies. To investigate the effects of Tau4RΔK and *para*^*bss*^ on circadian clock-driven neuronal computations in *Drosophila*, we asked how Tau4RΔK and *para*^*bss*^ modulate DN1p neuronal biophysical parameters related to neuronal activity. We recorded the spontaneous activity of the DN1p clock neurons at Zeitgeber time (ZT) 18-20 and compared the membrane potential dynamics with Tau4RΔK or the *para*^*bss*^ expression (**Fig. 1A**). We focused on ZT18-20 mid-night activity as our previous work demonstrated that DN1p activity during this timeframe exhibited regular firing patterns (Tabuchi et al., 2018), which turned irregular during mid-day recordings. We found that Tau4RΔK and *para*^*bss*^ induced remarkable alteration of the action potential initiation process in reduction of the instantaneous peak depolarization rapidness (dv/dt) during the depolarization phase (**Fig. 1B**). As we observed increased changes in the depolarization onset of action potentials in Tau4RΔK- or *para*^*bss*^-expressing DN1p circadian neurons, we hypothesized that Tau4RΔK or *para*^*bss*^ expression reduces the stability of spike onset properties. To quantitatively link the dynamical instability of spike onset phase kinetics to the formation of DN1p temporal spike patterns, we constructed a discrete-time Markov chain model (Nguyen et al., 2022). Individual DN1p action potentials were classified into three discrete states based on spike onset rapidness, meaning that each state represents a distinct regime of spike onset kinetics. We found that Markov chain analysis revealed that control DN1p neurons exhibited strong state persistence, remaining predominantly within a specific spike onset rapidness state (**Fig. 1C**). In contrast, Tau4RΔK- or *para*^*bss*^-expressing DN1p circadian neurons showed frequent transitions between states, indicating reduced stability of spike onset dynamics (**Fig. 1C**). The degree of dynamical instability is known to be quantified by the Lyapunov exponent (Fell et al., 1993; Ishizuka and Hayashi, 1996; O’Gorman et al., 2009). A Lyapunov exponent that is more positive suggests that the corresponding phase trajectories are diverging, and indicates the system is dynamically unstable (Fell et al., 1993; Ishizuka and Hayashi, 1996; Meng et al., 2022). Based on these perspectives, we reasoned that quantifications of Lyapunov exponent would be another metric to measure the dynamical instability in Tau4RΔK- or *para*^*bss*^-expressing DN1p circadian neurons in *Drosophila*. We found that higher Lyapunov exponent in Tau4RΔK- or *para*^*bss*^-expressing DN1p circadian neurons in *Drosophila*, compared to control (**Fig. 1D**). Visualization of the time-series variation of Lyapunov exponent with a discrete-time Markov chain (**Fig. 1E**) revealed that the dynamics were similar to those observed when the speed of action potentials was visualized. Our results suggest that visualization of Lyapunov exponent fluctuation via a discrete-time Markov chain could serve as a biophysical model of membrane dynamics variability. The time-series variation of Lyapunov exponent in control membrane potential showed strong state persistence in a specific class of Lyapunov exponent degree, whereas membrane potential in Tau4RΔK- or *para*^*bss*^-expressing DN1p circadian neurons showed frequent state transitions (**Fig. 1E**). To examine if this biophysical functional parameter was associated with macroscale variability of temporal spike patterns, we analyzed local variation (Holt et al., 1996) of the adjacent interspike interval from the same dataset and found that local variation in spike trains was also significantly increased in the DN1p clock neurons expressing Tau4RΔK or *para*^*bss*^ (**Fig. 1F**), suggesting that Tau4RΔK- and *para*^*bss*^-induced instability is associated with temporally proximate local operation of the DN1p clock neurons in *Drosophila*. Given that subtle alterations in the kinetics of membrane depolarization were associated with changes in the regularity of action potential firing patterns, we reasoned that these small-scale effects might propagate to influence global activity across broader brain regions. Accordingly, we examined local field potentials (LFP) recorded near pars intercerebralis (PI) neurons, the target of DN1p clock neuron projections (Barber et al., 2016; Chong et al., 2025), and quantified the signals using continuous wavelet transform. While power analysis of the raw LFP revealed no detectable differences (**Fig. 1G**), we hypothesized that hidden temporal structures might exist that are not readily apparent. To test this, we modeled the time series using an Ornstein–Uhlenbeck (OU) process (Lansky and Smith, 1989), and visualized their time-series process using the continuous wavelet transform (**Fig. 1H**). By quantifying the OU parameters such as time constant and noise amplitude (Ditlevsen and Lansky, 2005, 2006), we identified pronounced differences in LFP obtained from flies with Tau4RΔK- or *para*^*bss*^-expressing DN1p circadian neurons (**Fig. 1IJ**). The time constant was significantly larger in LFP recordings from flies expressing Tau4RΔK or *para*^*bss*^ in DN1p circadian neurons compared with controls, indicating reduced “instantaneity” of LFP dynamics (**Fig. 1I**). Furthermore, the noise amplitude was significantly elevated (**Fig. 1J**), reflecting increased fluctuations and instability. Finally, assuming that the presence of a unit root could contribute to the observed differences in the OU process, we performed an Augmented Dickey-Fuller (ADF) test to assess whether the time series are stationary (Jameson et al., 2024). A unit root indicates that a time series exhibits a stochastic trend, so a higher p-value in the ADF test reflects insufficient evidence to reject non-stationarity. We found that LFP recordings from flies expressing Tau4RΔK or *para*^*bss*^ in DN1p circadian neurons exhibited higher p-values (**Fig. 1K**), suggesting that their OU processes contain more non-stationary components.

**Fig. 1:**
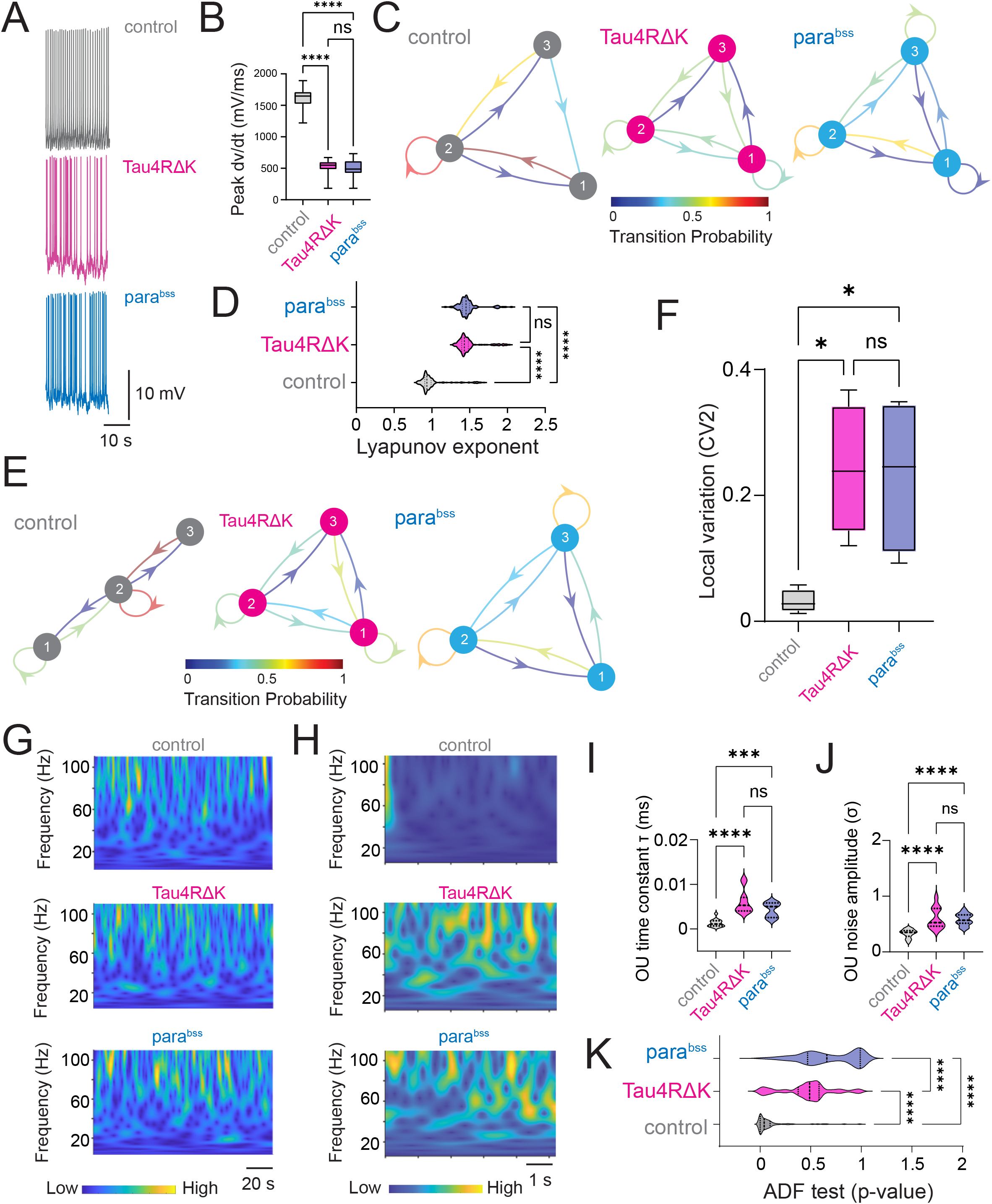
Tau4RΔK and *para*^*bss*^ induce dynamical instability of DN1p circadian neuronal activity at ZT18-20. (**A**) Membrane voltage traces in spontaneous activity in DN1p circadian neuron at ZT18-20 with/without Tau4RΔK or *para*^*bss*^ expression. (**B**) Quantification of peak speed of onset rapidness (dV/dt) of action potential of DN1p at ZT18-20 with/without Tau4RΔK or *para*^*bss*^ expression. (**C**) Discrete-time Markov chains visualizing stochastic process of state transition based on time series of individual action potential initiation process in DN1ps with/without Tau4RΔK or *para*^*bss*^ expression at ZT18-20. Categorization of the quantified maximum spike onset velocity (dV/dt) into three different classes based on their probability distribution. Class 1 was defined as the range from the minimum to the first quartile, class 2 was defined as the range from the first quartile to the third quartile, and class 3 was defined as the range from the third quartile to the maximum. The figure shows the transition matrix generated to calculate the probability of transitioning between these defined classes. Control shows state persistence, whereas Tau4RΔK or *para*^*bss*^ expression induces rapid state transition (**D**) Quantification of Lyapunov exponent on time series of membrane potential of DN1ps (**E**) Discrete-time Markov chains showing state transition based on time series of individual Lyapunov exponent (**F**) Spike variability quantify based on local variation between adjacent interspike intervals. Sample sizes: control: n=5, Tau4RΔK: n=4 *para*^*bss*^: n=4. (**G**) Continuous wavelet transform in time series of LFP recordings. intervals Sample sizes: control: n=8, Tau4RΔK: n=8 *para*^*bss*^: n=8. (**H**) Continuous wavelet transform in the time series of the Ornstein– Uhlenbeck (OU) model obtained from the LFP recordings. (**I**) OU time constant (τ) was defined as the exponential decay constant of the LFP autocorrelation. (**J**) OU noise amplitude (σ) was defined as the variability of LFP voltage fluctuations. (**K**) OU-modeled time series were analyzed by continuous wavelet transform, and their stationarity was assessed using Augmented Dickey–Fuller (ADF) tests. **p < 0.01, ****p < 0.0001. ns: non-significance based on one-way ANOVA followed by post-hoc Tukey tests.

### Antiepileptic Drugs Suppress Tau4RΔK- or *para*^*bss*^-induced Dynamical Instability

Previous studies have shown that antiepileptic drugs can restore hyperexcitability associated with Alzheimer’s disease (AD) pathogenesis (Sanchez et al., 2012; Tabuchi et al., 2015), beyond their canonical use in treating epileptic seizures. Thus, we decided to test if antiepileptic drug brivaracetam (BRV) can also restore Tau4RΔK- or *para*^*bss*^-induced dynamical instability in *Drosophila*. We found that administration of BRV suppressed the dynamical instability of spike onset rapidness in DN1p circadian neurons (**Fig. 2A**). To describe the stochastic process of the temporal structural state transition among distinct states of depolarization onset dynamics during action potential generation, we used a discrete-time Markov chain again. As we have already shown in **Fig. 1C**, the control had a strong recurrent preference in a preferred state, which generated a state persistence, which this trend was not observed in Tau4RΔK expression in DN1p neurons, and instead had more rapid state transition, but such altered state transitions of depolarization onset dynamics during action potential generation in time series were restored by administration of BRV (**Fig. 2B**). We also identified that increased local spike train variability found in Tau4RΔK or *para*^*bss*^-expressing DN1p was rescued by BRV (**Fig. 2C**). Distribution of interspike interval and instantaneous firing frequency altered in the DN1p clock neurons expressing Tau4RΔK or was also rescued by BRV (**Fig. S1**). Finally, we assessed the effectiveness of BRV in global brain states via the OU process (**Fig. 2D**) and found that BRV rescued both the elevated time constant (**Fig. 2E**) and the increased noise amplitude (**Fig. 2F**). Strikingly, evidence for the presence of a unit root was reduced, as reflected by the decreased p-value in the ADF test (**Fig. 2G**). These results suggest the increased dynamical instability induced by Tau4RΔK or *para*^*bss*^ has been restored by BRV administration in DN1p circadian neurons in *Drosophila*.

**Fig. 2:**
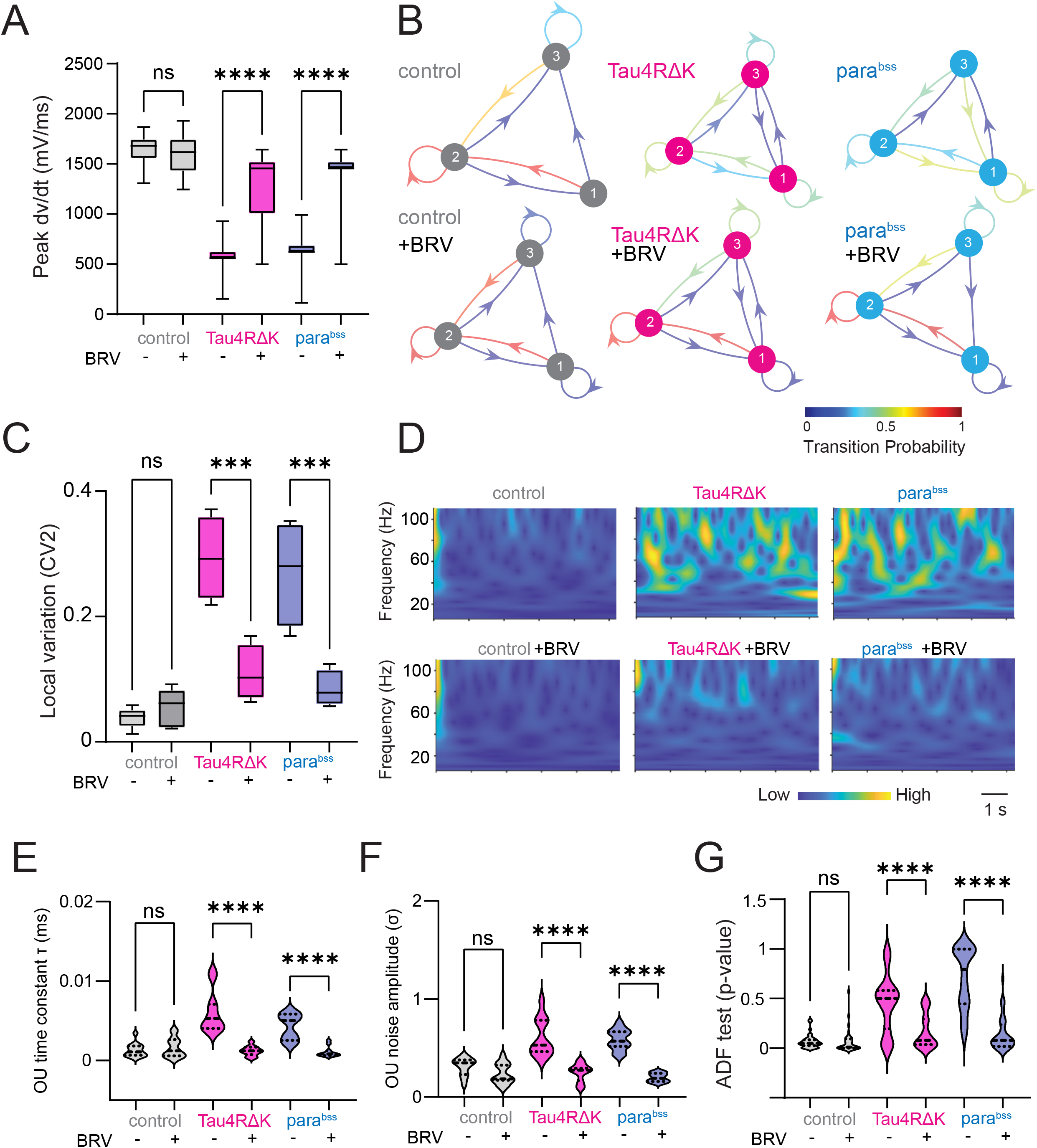
Antiepileptic drugs suppress instability in DN1p neural activity. (**A**) Quantification of peak speed of onset rapidness (dV/dt) of action potential of DN1p at ZT18-20 with/without Tau4RΔK or *para*^*bss*^ expression as well as the administration of the antiepileptic drug brivaracetam (BRV). (**B**) Discrete-time Markov chains visualizing stochastic process of state transition matrix based on time series of peak speed of onset rapidness of individual action potential initiation in DN1p with/without Tau4RΔK or *para*^*bss*^ expression as well as the administration of BRV. (**C**) Spike pattern variability based on local variation of interspike intervals, (**D**) Continuous wavelet transform in the time series of the Ornstein–Uhlenbeck (OU) model obtained from the LFP recordings. (**E**) OU time constant (τ) was defined as the exponential decay constant of the LFP autocorrelation. (**F**) OU noise amplitude (σ) was defined as the variability of LFP voltage fluctuations. (**G**) OU-modeled time series were analyzed by continuous wavelet transform, and their stationarity was assessed using Augmented Dickey–Fuller (ADF) tests. Sample sizes: control: n=8, control + BRVl: n=8, Tau4RΔK: n=5, Tau4RΔK + BRV: n=5, *para*^*bss*^: n=5, *para*^*bss*^ + BRV: n=5. **p < 0.01, ****p < 0.0001. ns: non-significance based on one-way ANOVA followed by post-hoc Tukey tests.

### Altered Sodium Current in DN1p Circadian Neurons Expressing Tau4RΔK or *para*^*bss*^

The voltage-gated sodium channel is one of the most important biophysical factors in shaping the membrane potential dynamics of neurons (Allen et al., 2017; Calvin, 1975), especially, initial depolarization during the action potential generation process (Do and Bean, 2003; Taddese and Bean, 2002). Given that *para*^*bss*^ is a mutation in the para sodium channel gene, and our observations of altered action potential initiation in both models, we examined if voltage-gated sodium channel currents are altered in DN1p circadian neurons expressing Tau4RΔK and *para*^*bss*^ at ZT18-20. We conducted voltage-clamp experiments using perforated-patch clamping, and pharmacological current isolation was performed to isolate sodium currents and remove all the other currents (See Methods). Because we found that changes in the initial depolarization during action potential generation influenced the variability of temporal spike patterns, we quantified fluctuations in the availability of voltage-gated sodium channels by measuring their inactivation state, which in turn affects subsequent spike patterns (Do and Bean, 2003; Taddese and Bean, 2002). Compared with controls **(Fig. 3A**), neither Tau4RΔK expression (**Fig. 3B**) nor *para*^*bss*^ (**Fig. 3C**) produced significant changes in the current– voltage relationship, as evidenced by quantification of the half-maximal voltage for voltage-dependent inactivation of voltage-gated sodium channels (**Fig. 3D**). This trend was also observed under BRV treatment (**Figs. 3E-H**). We next quantified the temporal kinetics of the non-stationary inactivation of these sodium currents using noise fluctuation analysis (**Figs. 3I–P**). Noise variability was calculated as the standard deviation of individual current traces subtracted from the mean trace (Sigworth, 1980). This analysis revealed that noise variability was significantly increased by both Tau4RΔK (**Fig. 3J**) and *para*^*bss*^(**Fig. 3K**), compared to control (**Fig. 3I**). Among these comparisons, Tau4RΔK showed the highest noise variability, surpassing that of *para*^*bss*^(**Fig. 3L**), suggesting a strong disruption of sodium channel inactivation. Remarkably, the increased noise variability induced by Tau4RΔK and *para*^*bss*^ was restored by BRV treatment (**Figs. 3M–P**), suggesting that BRV effectively stabilizes sodium channel inactivation and thereby normalizes spike pattern variability.

**Fig. 3:**
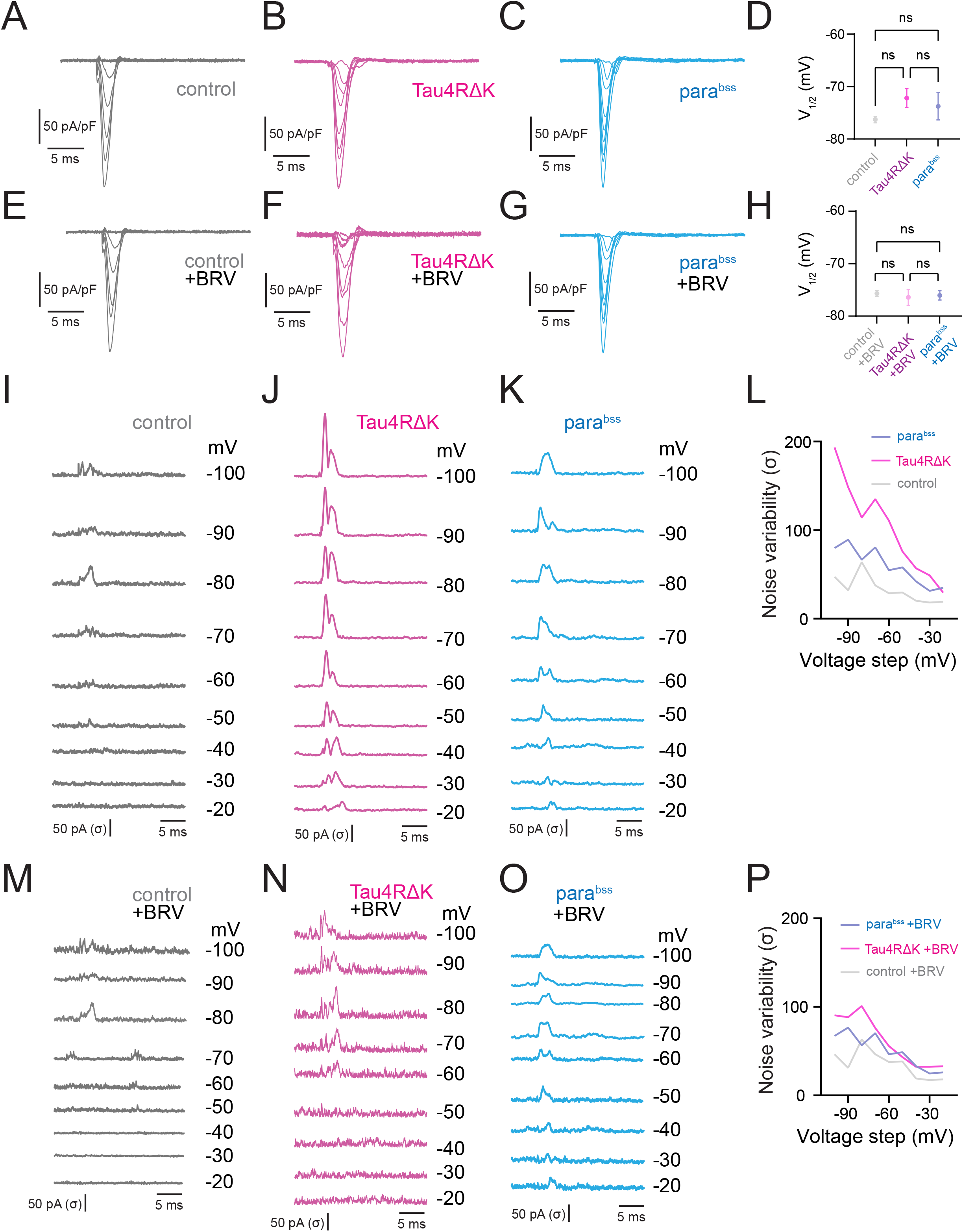
Tau4RΔK and *para*^*bss*^ alter biophysical properties of sodium currents in DN1ps. Averaged sodium current traces in DN1p circadian neurons: control (**A**), Tau4RΔK expression (**B**), *para*^*bss*^ expression (**C**), and their half-maximal voltage quantification (**D**). Traces under BRV administration: control + BRV (**E**), Tau4RΔK + BRV (**F**), *para*^*bss*^ + BRV (**G**), and corresponding half-maximal voltage quantification (**H**). Currents were measured during a 0 mV test pulse preceding step depolarization from –100 to –20 mV (steady-state inactivation protocol). (**I–K**) Temporal fluctuations of sodium currents were visualized as the standard deviation of individual current traces subtracted from the average: control (**I**), Tau4RΔK (**J**), *para*^*bss*^ (**K**). (**L**) Maximum noise variability during the non-stationary inactivation process across pre-pulse depolarization from–100 to –20 mV. (**M–O**) Same as (**I–K**) after BRV treatment: control + BRV (**M**), Tau4RΔK + BRV (**N**), *para*^*bss*^ + BRV (**O**), with maximum noise variability quantified in (**P**). Sample sizes: control, n = 6; control + BRV, n = 6; Tau4RΔK, n = 4; Tau4RΔK + BRV, n = 5; parabss, n = 5; parabss + BRV, n = 5. Statistical significance: *p < 0.05, **p < 0.01, ***p < 0.001, ****p < 0.0001; ns, not significant (one-way ANOVA with post-hoc Tukey tests).

### Increased local variance of spike irregularity in AD and epilepsy patient iPSC-derived neurons

We wish to test whether phenotypes that have been observed in fly AD model can also be observed in human AD and epilepsy. Since patient iPSC-derived neurons in conjunction with *Drosophila* models offer a powerful genetic approach for advancing our understanding of neurological disorders (Vadodaria et al., 2020), we utilize human iPSC-derived neuronal cultures obtained from AD and epilepsy patients. Electrophysiological characterizations of human iPSC-derived neuronal networks using microelectrode arrays and their disease relevance have already been established (Gunhanlar et al., 2018; Halliwell et al., 2021; Page et al., 2022), and thus we applied this quantitative approach to test if AD and epilepsy patient iPSC-derived neurons show similar phenotypes observed in *Drosophila* models. Spontaneous action potential firing patterns were extracellularly identified from healthy control, AD patients, and epilepsy patients, as well as applications of BRV (**Fig. 4A and Fig. S2**). Although the distributions of interspike intervals (**Fig. S2A**) and instantaneous spike frequency (**Fig. S2B**) were comparable, the mean firing rate differed between control and BRV-treated controls, as well as between control and epilepsy patient neurons (**Fig. 4B**). More strikingly, local spike train variability was significantly elevated in both AD and epilepsy patient iPSC-derived neurons (**Fig. 4C**), with epilepsy neurons exhibiting the highest variability. BRV treatment rescued this increased variability in both disease groups (**Fig. 4C**), mirroring observations in *Drosophila* models. Interestingly, epilepsy patient neurons displayed an intermediate level of variability between control and AD, suggesting a spectrum of neuronal instability across neurological conditions. We sought to address the seemingly contradictory situation where no change is observed in the distribution structure of interspike interval histograms and instantaneous spike frequencies (**Fig. S2**), yet local spike variability is observed (**Fig. 4C**). We used neuroinformatics approaches to characterize latent features that cannot be manifested in distribution structure of spike sequence. Specifically, we performed Gaussian mixture model (GMM) clustering and random forest classification. In the GMM clustering, as previously demonstrated (Tabuchi et al., 2018), we focused on adjacent interspike interval structure, as the difference was observed in local variability (**Fig. 4C**). Control iPSC-derived neurons exhibited clearly separable, binarized emergence of highly patterned spike chains, whereas AD and epilepsy patient neurons showed disrupted GMM cluster probabilities (**Fig. 4D**). Importantly, BRV treatment restored the binarized emergence of highly patterned spike chains in both AD and epilepsy neurons. We next performed random forest classification to quantify probabilistic similarity among the datasets, by building a multitude of decision trees at training time and combining their predictions to make more accurate and robust classifications. Time series of interspike interval and instantaneous spike frequencies were used as target variable and predictor variables. Classification results were similar for both interspike interval and instantaneous spike variables (**Fig. 4E**). The probabilistic distance analysis revealed a hierarchical relationship: control neurons were most distinct from epilepsy neurons, with AD neurons occupying an intermediate position in the “No drug” condition. This suggests that AD represents a partial phenocopy of epilepsy at the neuronal dynamics level. Compared to “No drug” conditions, the probabilistic distance between control and both disease groups was significantly reduced in the presence of BRV. Both AD and epilepsy patient neurons exhibit similar dynamic instabilities, differing in magnitude, and this suggests that the spectrum of neuronal instability may underlie distinct manifestations, from episodic seizures to progressive neurodegeneration. Notably, antiepileptic drug BRV successfully rescued these instabilities in both conditions, highlighting microscale neuronal dynamics as a potential therapeutic target across neurological disorders.

**Fig. 4:**
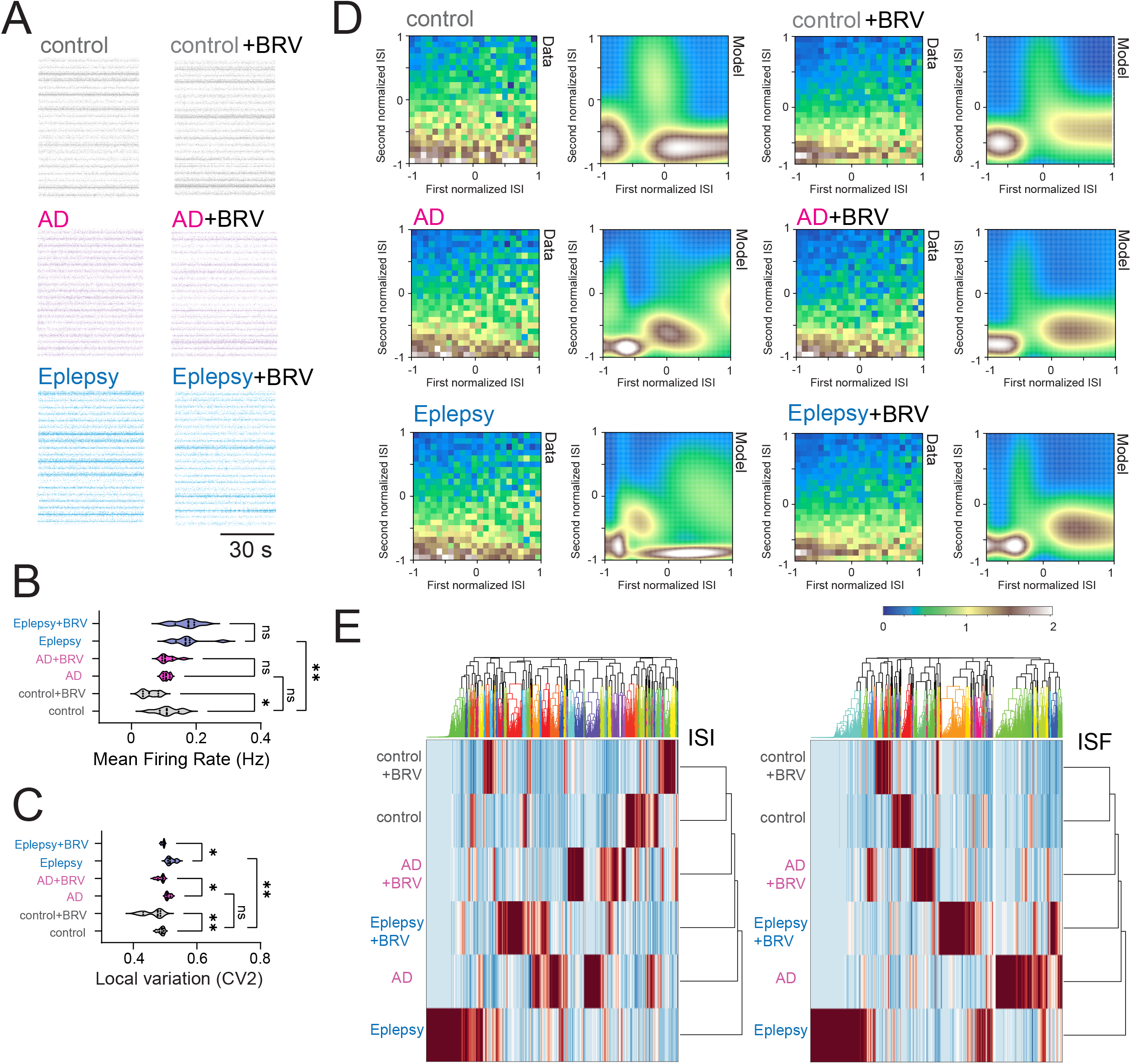
Local spike irregularity in spontaneous activity of human iPSC-derived neuronal culture obtained from AD and epilepsy patients. (**A**) Raster plot visualizing neural spike trains. (**B**) Quantification of mean firing rate, (**C**) quantification of local spike pattern variability based on adjacent interspike intervals of human iPSC-derived neuronal culture at DIV 55. (**D**) Structural motif in spontaneous activity of human iPSC-derived neuronal culture obtained from AD and epilepsy patients, showing distinctive profiling in emergence probability of adjacent interspike interval distribution visualized by GMM clustering. (**E**) Hierarchical clustering analysis of temporal sequence of interspike interval and hierarchical clustering analysis of temporal sequence of instantaneous spike frequency. 2 healthy controls (CW50065 and CW50023), 3 AD patients (CW50174, CW50114, and CW50137), and 3 epilepsy patients (CW60236, CW60231, and CW60130). Sample sizes: n=320 from 20 cultures for each condition. Statistical significance: *p < 0.05, **p < 0.01, ***p < 0.001, ****p < 0.0001; ns, not significant (one-way ANOVA with post-hoc Tukey tests).

## Discussion

Our study supports a role for dynamical instability of the microscale action potential initiation process in the production of local spike variability that causes aberrant macroscale behaviors mediated by Tau4RΔK and *para*^*bss*^ in the circadian network, which conceptually providing evidence that the usually neglected subtle microscale biophysical dynamical instability plays a key role in regulating the macroscale dynamics of neuronal activity, sleep, and longevity in *Drosophila*. Furthermore, the observed correlation between fly- and iPSC-derived neurons underscores the translational significance of our findings, suggesting that subtle microscale biophysical changes can manifest as overt macroscale pathologies that can be shared between AD and epilepsy (Minkeviciene et al., 2009). In this study, we leveraged both *Drosophila* and human iPSC-derived neuron models to understand how these alterations drive disease progression, how they contribute to global network dysfunction, and how they might be therapeutically targeted. AD and epilepsy both exhibit significant disruptions in circadian control of sleep architecture, and therefore it is possible that shared properties between AD and epilepsy can be explained by sleep disturbance (Hanke et al., 2022). It is well known that AD and epilepsy display circadian dysregulation mediated through pathological changes to circuits that regulate the suprachiasmatic nucleus (Gerstner et al., 2014; Paul et al., 2018; Sanabria et al., 1996). In AD, these disruptions manifest in symptoms such as “sundowning,” while in epilepsy, seizures often occur in relation to distinct phases of the sleep-wake cycle (Gerstner et al., 2014; Karoly et al., 2018; Vitiello et al., 1992). These observations underscore the importance of sleep disturbances in neurodegenerative conditions and epilepsy.

In fly neurons, we found increased variability in voltage-gated sodium currents during non-stationary inactivation, suggesting a candidate ionic contributor to altered spike timing. Antiepileptic treatment reduces sodium-current variability and stabilizes spike initiation in the fly models, while producing a parallel improvement in spike-timing instability in patient-derived human neurons. These findings are consistent with a model in which fluctuations in sodium-channel availability during non-stationary inactivation increase trial-to-trial variability in spike initiation. Once a channel inactivates, the subsequent action potential timing is determined by the fraction of sodium channels that recover to a functional state (Armstrong, 2006). Decreased cooperativity means fewer sodium channels need to open to initiate an action potential, increasing the variability (or “noise”) of the process (Naundorf et al., 2006). This heightened noise can accumulate over time, resulting in local spike train variability and erratic firing patterns (Kuriscak et al., 2012). In this view, altered mean excitability is not the only relevant dimension of dysfunction; variability itself may represent a disease-relevant physiological feature.

In the fly models, unstable spike initiation is also associated with altered circuit- and brain-state readouts. The importance of finding subtle biophysical phenotypes is underscored by their downstream consequences for larger-scale brain functions such as sleep regulation and memory consolidation (Hanke et al., 2022). Although our data do not establish a complete causal chain from channel-level variability to organism-level pathology, they support the idea that subtle cellular instability can be reflected in higher-order network dynamics. This interpretation is compatible with the general principle that small fluctuations in spike timing may be amplified by recurrent interactions, especially in circuits that depend on precise timing and state stability.

The convergence between *Drosophila* and patient-derived human neurons should likewise be interpreted cautiously. We do not infer a one-to-one translation between these preparations or claim a universal mechanism for AD and epilepsy. Rather, the cross-model agreement identifies unstable spike initiation as a conserved phenotype that recurs across distinct genetic and disease contexts. This shared phenotype provides a useful intermediate level of description between molecular perturbation and systems dysfunction.

The pharmacological results further support this framework. In fly neurons, antiepileptic drugs reduce sodium-current variability together with spike-timing instability, and human neurons show a parallel improvement in firing stability. The precise neuroprotective mechanisms of antiepileptic drugs have remained incompletely understood (Klitgaard et al., 2016; Steinhoff and Staack, 2019), but our findings are consistent with a functional contribution of sodium-channel-dependent variability to the phenotype, although they do not by themselves prove that this is the only mechanism involved. More broadly, they suggest that stabilizing intrinsic neuronal dynamics may complement therapeutic strategies aimed solely at shifting mean excitability

Several limitations should be noted. The *Drosophila* recordings were obtained from a specialized neuronal population, and the extent to which the same dynamical phenotype generalizes to other cell types remains unknown. Human iPSC-derived neurons likewise lack the full circuit, glial and metabolic context of the intact brain. In addition, both AD and epilepsy are heterogeneous disorders, so sodium-current variability is unlikely to account for all forms of dysfunction across patients or models. With these caveats, the present study provides a tractable starting point for multiscale analysis. By defining unstable spike initiation as a measurable and reversible phenotype, and by nominating sodium-current variability as one candidate contributor, our work offers a framework for testing how subtle biophysical perturbations in intrinsic excitability may scale toward circuit dysfunction in neurological disease.

## Materials and Methods

### *Drosophila* genetics

All lines were obtained from the Bloomington *Drosophila* Stock Center (Bloomington, IN, USA), except for *UAS-Tau4RΔK*. To target and visualize DN1p neurons, *R18H11-Gal4* line (BDSC: 48832) and *UAS-CD4-tdGFP* line (BDSC: 35836) were used. *UAS-Tau4RΔK* and *UAS-para*^*bss*^ transgenic flies were generated in the iso31 background using standard techniques (Rainbow Transgenics) from the Tau4RΔK sequence obtained from Dr. Philip Wong (Li et al., 2016) and para^bss^ sequence (Parker et al., 2011) was fully synthesized.

Standard genetic recombination techniques were employed to generate flies expressing multiple transgenes. Flies were fed standard *Drosophila* food containing molasses, cornmeal, and yeast. They were housed in a 25°C incubator (DR-36VL, Percival Scientific, Perry, IA, United States) under 12 h:12 h light-dark cycles and 65% humidity. Female flies were used for all experiments. All experiments were performed at 25 °C and in compliance with all relevant ethical regulations for animal testing and research at Case Western Reserve University.

### Human iPSC-Derived Neurons

Excitatory neurons differentiated from cryopreserved human induced pluripotent stem cells (hiPSCs) were cultured and used for all experiments. We obtained hiPSCs from Elixirgen Scientific (Elixirgen Scientific, Inc., Baltimore, MD, USA), derived from five individuals: CW50023 (69-year-old man, Caucasian, not Latino, healthy), CW50065 (74-year-old woman, Caucasian, not Latino, healthy), CW50174 (76-year-old woman, Caucasian, not Latino, Alzheimer’s diagnosis), CW50114 (72-year-old woman, Caucasian, not Latino, Alzheimer’s diagnosis), and CW50137 (70-year-old woman, Caucasian, not Latino, Alzheimer’s diagnosis). We also used additional hiPSC-derived neuronal cultures obtained from epilepsy patients through Elixirgen Scientific’s Quick-Neuron platform (Lu et al., 2023; Yuan et al., 2020). Epilepsy lines were derived from individuals CW60236, CW60231, and CW60130, all of whom experienced grand mal and/or absence seizures. Despite the heterogeneity in seizure phenotypes, these lines were considered collectively as the “epilepsy” group. The healthy donor line, which exhibited no seizure activity, served as the control. All hiPSC lines were differentiated into excitatory neurons prior to use. For the first 10 days in vitro (DIV), cells were maintained in Elixirgen Scientific’s neuronal maintenance medium. Beginning at DIV 11, cultures were transitioned to Neurobasal Plus medium supplemented with B-27 Plus (Thermo Fisher). All cultures were maintained on 8×8 electrode arrays with 20-µm electrodes, compatible with the MED64 multi-array system amplifier.These hiPSCs were distributed by the California Institute of Regenerative Medicine (CIRM) Human Induced Pluripotent Stem Cell Repository, and differentiated using proprietary transcription factor-based methods. The Quick-Neuron™ EX-SeV kit (Elixirgen Scientific, Inc.) induced differentiation of the cells into excitatory neurons for the ten days leading to plating. Cell plates with MED 8×8 Probe 20 μm electrodes, 100 μm distance, and 10 mm height were utilized (P210A or R210A, AutoMate Scientific, Inc., CA, USA). The plates were initially coated with 0.01% Poly-L-ornithine (P4957, sigma) diluted in PBS, and allowed to incubate at 37°C overnight. The following day the same plates were washed with PBS, coated with 10 μg/mL laminin in PBS (#23017015, Thermo Fisher Scientific), and allowed to incubate overnight at 37°C. The hiPSC neurons were plated into each well at an approximate density of 10,000 cells/plate. Upon plating, the hiPSCs were maintained with Medium iN(G2P) (Elixirgen Scientific, Inc.). After the initial day of plating, Medium N(G2P) (Elixirgen Scientific, Inc.) was used to maintain the cells for another 5 weeks. The medium was regularly (every 3-4 days) changed by pipetting out approximately 50% of the medium, and replacing this volume with fresh medium. Electrophysiological recordings from the hiPSC neurons were performed by using MED64 Multi-electrode Array System amplifier (MED-A16HM1, AutoMate Scientific) with a dedicated temperature controller (MED-CP04, AutoMate Scientific). The data were sampled at 20 kHz with the interface (National Instruments), which was controlled on a computer using MED64 Mobius Software (MED-MS64AT03, AutoMate Scientific). The recording was conducted in DIV 55.

### Antiepileptic drugs

Brivaracetam (BRV, B677645, Toronto Research Chemicals) were used as antiepileptic drugs. BRV was prepared as a 94.2 mM stock solution dissolved in dimethyl sulfoxide, and was used to prepare for a final concentration of 20 μM with water containing 5% sucrose and 2% *Drosophila* agar medium and filled with 5 mm depth into the glass tubes of 5 mm diameter x 65 mm length. Flies (0-1 days old) that were collected were fed these antiepileptic drugs in prepared glass tubes for 48 hours in a 25°C incubator under 12 h:12 h light-dark cycles and 65% humidity. After this preincubation, flies (2-3 days old after the preincubation) were used for experiments.

### Electrophysiological Recordings

We conducted electrophysiological recordings from DN1p neurons with *ex vivo* configuration (i.e., isolated brain preparation). All experiments were performed at ZT18-20. Flies (2-3 days old) were chilled on ice (up to 10 min) for anesthesia and placed in a dissecting chamber following isolation of the head. Brains were removed and dissected in a *Drosophila* physiological saline solution (101 mM NaCl, 3 mM KCl, 1 mM CaCl_2_, 4 mM MgCl_2_, 1.25mM NaH_2_PO_4_, 20.7 mM NaHCO_3_, and 5 mM glucose; with osmolarity adjusted to 235-245 mOsm and pH 7.2), which was pre-bubbled with 95% O_2_ and 5% CO_2_. The glial sheath surrounding the brain was focally and carefully removed by using sharp forceps after treating with an enzymatic cocktail, collagenase (0.1 mg/mL), protease XIV (0.2 mg/mL), and dispase (0.3 mg/mL), at 22°C for 1 min to increase the likelihood of a successful recording. Using a small stream of saline, which was pressure-ejected from a large-diameter pipette, the surface of the cell body was cleaned under a dissection microscope. DN1p neurons were visualized via tdGFP fluorescence by using a PE300 CoolLED illumination system (CoolLED Ltd., Andover, UK) on a fixed-stage upright microscope (BX51WI; Olympus, Japan). One neuron per brain was recorded.

### Intracellular recordings

Sharp electrode intracellular recordings of DN1p neurons were performed by using sharp electrodes having “membrane-coating” (Jameson et al., 2024). The purpose of having “membrane-coating” is to obtain stable and low-noise performance in the electrodes (Jameson et al., 2024). Lipid preparation involved hour-long ultrasonication of egg lecithin (Sigma 440154, 7.6 g/L) plus cholesterol (Sigma C8667, 2 mM) within hexane-acetone medium at room temperature. Solvent removal occurred via nitrogen flow followed by vacuum exposure. The dried lipids were reconstituted in paraffin oil mixed with squalene (70:30 v/v) and equilibrated at 80°C overnight. Coating was achieved by “tip-dip” method based on the interface between the water and oil phases(Jameson et al., 2024). The sharp electrode was made from quartz glass with a filament (OD/ID: 1.2/0.6mm) with a laser-based micropipette puller (P-2000, Sutter instrument) and backfilled with 1 M KCl, with resistances of 120–190 MΩ. Solutions were filtered by using a syringe filter with a pore size of 0.02 μm (Anotop 10, Whatman). We inserted the electrode into the region having dense tdGFP signals near the cell body. We were unsure whether this region is corresponding to the Distal Axonal Segment (Ravenscroft et al., 2020), which is thought to be a spike generation site having dense expression of voltage-gated sodium channels. However, we could record a bigger amplitude of action potential from this region, compared to the cell body. During the insertion process of the electrode, tdGFP signals were used for initial visual inspection, and the depth direction of insertion was determined by change of sound (Model 3300 Audio Monitor, A-M Systems) and generation of membrane potential. This process was facilitated by adding “buzz” pulses at the moment just before the electrode is about to cross the membrane. The duration of pulse was determined by “advance air shooting”. If the tip of the electrode can be visually seen physically shaking, the duration was determined to be too long. We utilized the longest duration in the range that the electrodes do not move, which was typically between 2-5 ms. Recordings of membrane potentials commenced after membrane potential was stable, as it usually takes at several minutes to stabilize the cell membrane potential. Recordings were acquired with an Axoclamp 2B with HS-2A x 1 LU headstage (Molecular Devices), and sampled with Digidata 1550B interface, which were controlled on a computer using pCLAMP 11 software. The signals were sampled at 10 kHz and low-pass filtered at 1 kHz.

### Perforated patch-clamp recordings

Perforated patch-clamp of DN1p neurons were performed from somatic recording. Patch pipettes (9-12 MΩ) for perforated patch-clamp were constructed from borosilicate glass capillary (without filament) using a Flaming-Brown puller (P-97, Sutter Instrument) and further polished with a MF200 microforge (WPI) prior to filling internal pipette solution (102 mM cesium chloride, 0.085 mM CaCl_2_, 0.94 mM EGTA, 8.5 mM HEPES, 4 mM Mg-ATP, 0.5mM Na-GTP, 17 mM NaCl; pH7.2). Patch pipettes were also coated with membrane to achieve for stable and low-noise performance (Jameson et al., 2024). Escin was prepared as a 50 mM stock solution in water (stored up to 2 weeks at −20°C) and was added fresh into the internal pipette solution to a final concentration of 50 μM. Filling syringes were wrapped with aluminum foil, owing to the light-sensitivity of escin. Pipette tips were briefly dipped for 1 s or less into a small container with escin-free internal pipette solution, and then were back-filled with the escin-containing solution from the filling syringe. Air bubbles were removed by gentle tapping. Escin pipette solutions remained stable for several hours after mixing in the filling syringe, with no evidence of precipitate formation. Junction potentials were nullified prior to high-resistance seal formation. After the high-resistance seal formation, perforated patches were allowed to develop spontaneously over time. After breakthrough became evident, as determined by the gradual development of a large capacitance transient in the seal test window, access resistance monitoring was initiated employing the membrane test function. After that point, access resistance was monitored continuously during the final completion of perforation process, until it becomes stable (access resistance stably < 40 MΩ). Cells that showed signs of “mechanical” break-in (as determined by measured by a significant increase of time constant of transient capacitive current) were excluded for further data acquisition. During the recording, the bath solution was continuously perfused with saline by means of a gravity-driven system. To isolate voltage-dependent sodium currents during their inactivation process, we perform voltage-clamp recordings with constant test pulse of 0 mV before the step depolarization from -100 mV to -20mV with 10 mV increment, loaded with a cesium chloride patch pipette internal solution and in a bath solution containing 0.001 mM cadmium chloride (CdCl_2_), 0.005 mM M 4-Aminopyridine (4-AP), and 0.02 mM Tetraethylammonium (TEA).

Recordings were acquired with an Axopatch 1D, 200A or 200B amplifier (Molecular Devices) and sampled with Digidata 1550B (Molecular Devices) controlled by pCLAMP 11 (Molecular Devices). The voltage signals were sampled at 20 kHz and low-pass filtered at 1 kHz.

### LFP recordings

Local field potential (LFP) recordings were obtained *in vivo* from regions adjacent to pars intercerebralis (PI) neurons using a large-diameter borosilicate glass pipette electrode (resistance <1 MΩ). In vivo electrophysiological preparation of PI neurons was performed as previously described(Chong et al., 2025; Tabuchi et al., 2018) ; briefly, flies were anesthetized by chilling for 10 min and secured to a 0.025 mm-thick metal shim using dental wax or UV-curable adhesive. The cuticle was carefully removed to expose the brain surface, and the tethered fly was mounted in a recording chamber filled with Drosophila physiological saline. Signals were amplified using two battery-powered DAM50 amplifiers (World Precision Instruments) and digitized via a PCIe-6341 data acquisition interface (National Instruments) at a sampling rate of 10 kHz, followed by low-pass filtering at 1 kHz.

### Analysis of electrophysiology data (*Drosophila*)

Electrophysiological analysis was performed in MATLAB (MathWorks). Basic spike statistics such as mean firing rate, interspike interval histogram, instantaneous firing rate, and spike variability were quantified as describe previously. To estimate the temporal structure of state transition of spike waveform in spike sequences, we used discrete-time Markov chain. We first classified quantified maximum spike onset rapidness (dV/dt) into three groups based on their probability distribution. The range from minimum to the first quartile was defined as class 1, the range from the first quartile to the third quartile was defined as class 2, and the range from the third quartile to maximum was defined as class 3. We then created a transition matrix to calculate transition probability.

To quantify the dynamical instability of membrane potential, we used the Lyapunov exponent. The Lyapunov exponent was computed as follows.

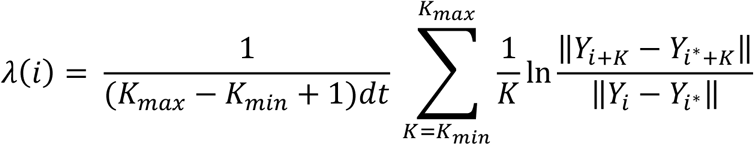

where *K_max_* and K_min_ represent the minimum and maximum value of expansion range of trajectories projected in phase space of the time variation of the membrane potential and the value of the time derivative of the time variation of the membrane potential, and dt represents sampling time of the trajectories. A selected trajectory Yi as a reference trajectory and the rate of expansion of the distance between the reference orbit and another orbit Yi+K from an initial condition that is only an instantaneously small distance* away from the reference orbit is exponential, and the limit value of this exponential is the Lyapunov exponent, which indicates an exponential increase in the distance between the orbits, so that a Lyapunov exponent Therefore, a large Lyapunov exponent indicates a high degree of instability in the dynamical system, which is membrane potential dynamics of the DN1ps neurons in the present study.

An Ornstein–Uhlenbeck (OU) model was constructed using parameters extracted from LFP recordings. LFP signals were first preprocessed and mean-centered. The time constant of the OU process was estimated from the temporal autocorrelation structure of the LFP by fitting an exponential decay function to the autocorrelation function, corresponding to the inverse of the mean-reversion rate. The noise amplitude was quantified from the variability of the LFP signal, defined as the standard deviation of voltage fluctuations after detrending. Numerical simulations were implemented using the Brian simulator (Tabuchi et al., 2018) with a fixed time step matching the original LFP sampling interval. The simulated OU time series were subsequently analyzed using continuous wavelet transform to decompose the signals into frequency-specific components. For each frequency band, the resulting wavelet-transformed time series was subjected to an Augmented Dickey-Fuller (ADF) test to assess stationarity (Jameson et al., 2024). The ADF test evaluates the null hypothesis that a time series contains a unit root, which indicates non-stationarity characterized by a stochastic trend. Thus, higher p-values correspond to insufficient evidence to reject the presence of a unit root. Differences in p-value distributions across genotypes were interpreted as reflecting differences in the degree of non-stationary dynamics embedded in the OU-modeled LFP signals.

Steady-state inactivation of voltage-dependent sodium currents were quantified by constant test pulse of 0 mV before the step depolarization from -100 mV to -20mV with 10 mV increment. The nonstationary noise analysis of voltage-gated sodium channels during their inactivation process was processed based on a previous study (Sigworth, 1980). Specifically, we defined the variability of the nonstationary noise as the standard deviation calculated from the residual fluctuations in each data set after subtracting the time series of the averaged current. The larger deviations are near the time of the peak of the current. The variance was assumed to be primarily due to two independent sources of current fluctuations: the stochastic gating of sodium channels and the thermal noise background in the voltage clamp.

### Analysis of electrophysiology data (Human iPSC-Derived Neurons)

In contrast to intracellular recordings from *Drosophila*, electrophysiology data of human iPSC-derived neurons were extracellularly measured. Thus, we focus on temporal spike sequence analysis, rather than membrane potential dynamics. Action potential detection was based on voltage peaks deriving from capacitive currents. Baseline adjustment was performed, and action potential was registered based on manually determined threshold. For GMM model, adjacent interspike interval probability was used as a parameter, and the number of Gaussian components for data fitting was manually determined by visual inspection. For Random Forest Classification, temporal order of individual interspike interval and instantaneous spike frequency was used as dataset, and bootstrapped sampling method was implemented. Decision trees were constructed using the bootstrapped dataset, and the individual tree predictions were averaged to obtain the final prediction.

### Statistical Analysis

All statistical analyses were done using Prism software (GraphPad). For comparisons of 2 groups of normally or non-normally distributed data, t tests or Mann-Whitney U-tests were performed, respectively. For more than 2 multiple group comparisons, one-way ANOVA followed by post-hoc Tukey tests was used. A p-value < 0.05 is considered a statistically significant test result. Asterisks indicate p-values, where ∗p < 0.05, ∗∗p < 0.01, ∗∗∗p < 0.001, and ****p < 0.0001. ns indicates a non-significant test result. Error bars in box plots are minimum and maximum, and box represents interquartile range. Error bars in all bar graphs are means ± SEM averaged across experiments, but error bars in non-bar graphs are means SD. Violin plots are based on kernel density estimation implemented in Prism software.

## Acknowledgments

We thank Keisuke Sakurai for the loan of electrophysiological equipment and Joseph Monaco for technical advice on computational modeling. We also thank Ben Strowbridge, Heather Broihier, and Dominique Durand, along with members of the Tabuchi lab for discussion, and the Light Microscopy Imaging Core at Case Western Reserve University for help with confocal microscopy. Funding: This work was supported by grants from the National Institutes of Health (R00NS101065 and R35GM142490), Whitehall Foundation, BrightFocus Foundation (A2021043S), PRESTO grant from Japan Science and Technology Agency (JPMJPR2386), Research Corporation for Science Advancement (SA-MBC-2024-080c), and the Tomizawa Jun-ichi and Keiko Fund of the Molecular Biology Society of Japan for Young Scientists.

## Supplementary Figures

**Fig S1:** Additional electrophysiological analysis for the data used in Figs.1 and 2.

(**A**) Probabilistic distribution structure of interspike intervals distribution and (**B**) probabilistic distribution structure of instantaneous spike frequency obtained from spontaneous firing in DN1p circadian neurons with/without Tau4RΔK or *para*^*bss*^ expression.

**Fig S2:** Additional electrophysiological analysis for the data used in Fig.4.

(**A**) Probabilistic distribution structure of interspike intervals distribution and (**B**) probabilistic distribution structure of instantaneous spike frequency obtained from spontaneous firing in human iPSC-derived neuronal culture obtained from AD and epilepsy patients.

