## Supplementary figures and images for "Neuronal microscale biophysical instability mediates macroscale network dynamics shaping pathological manifestations"

### Supplemental Data 1

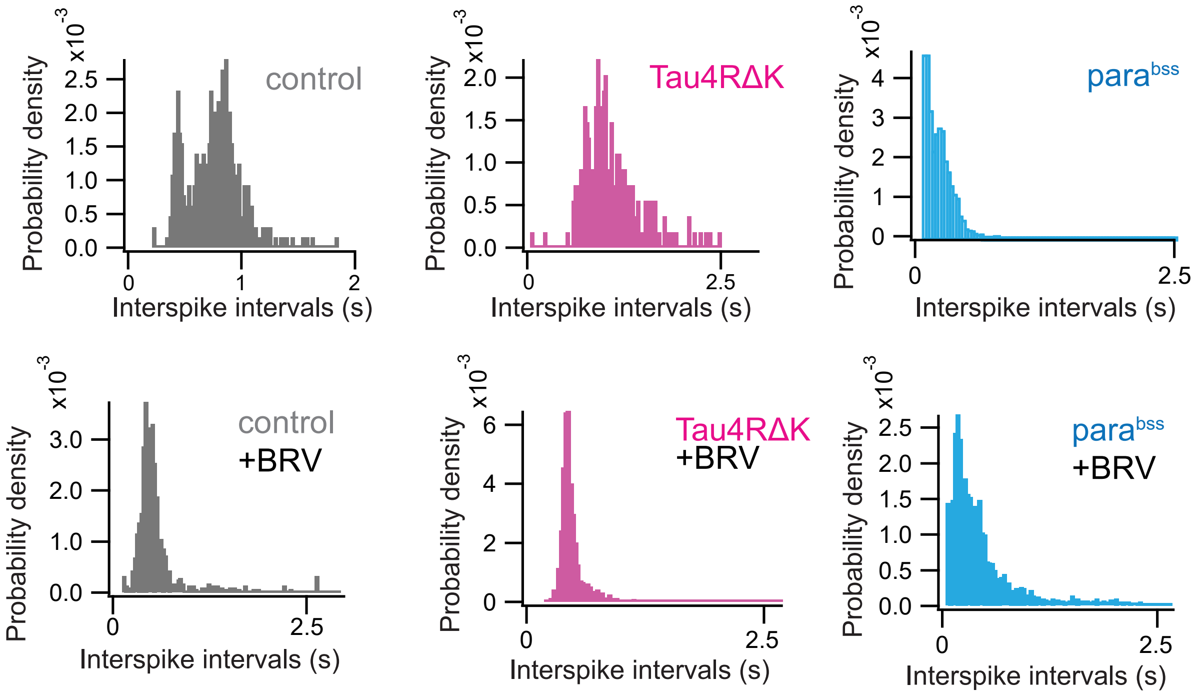

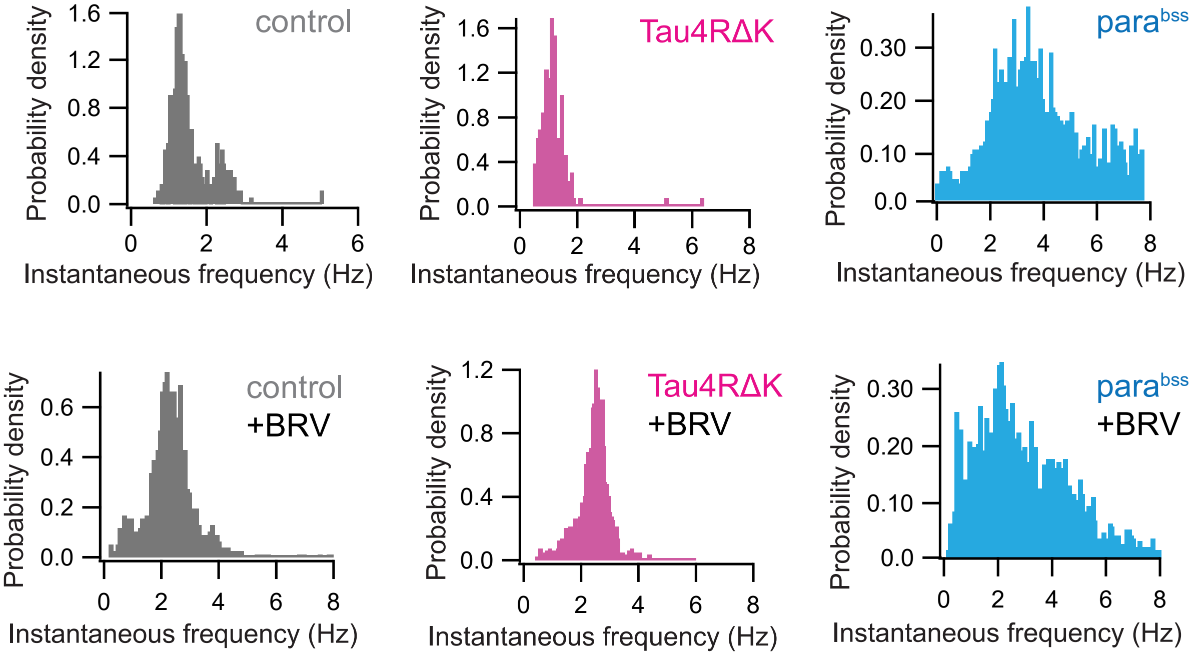

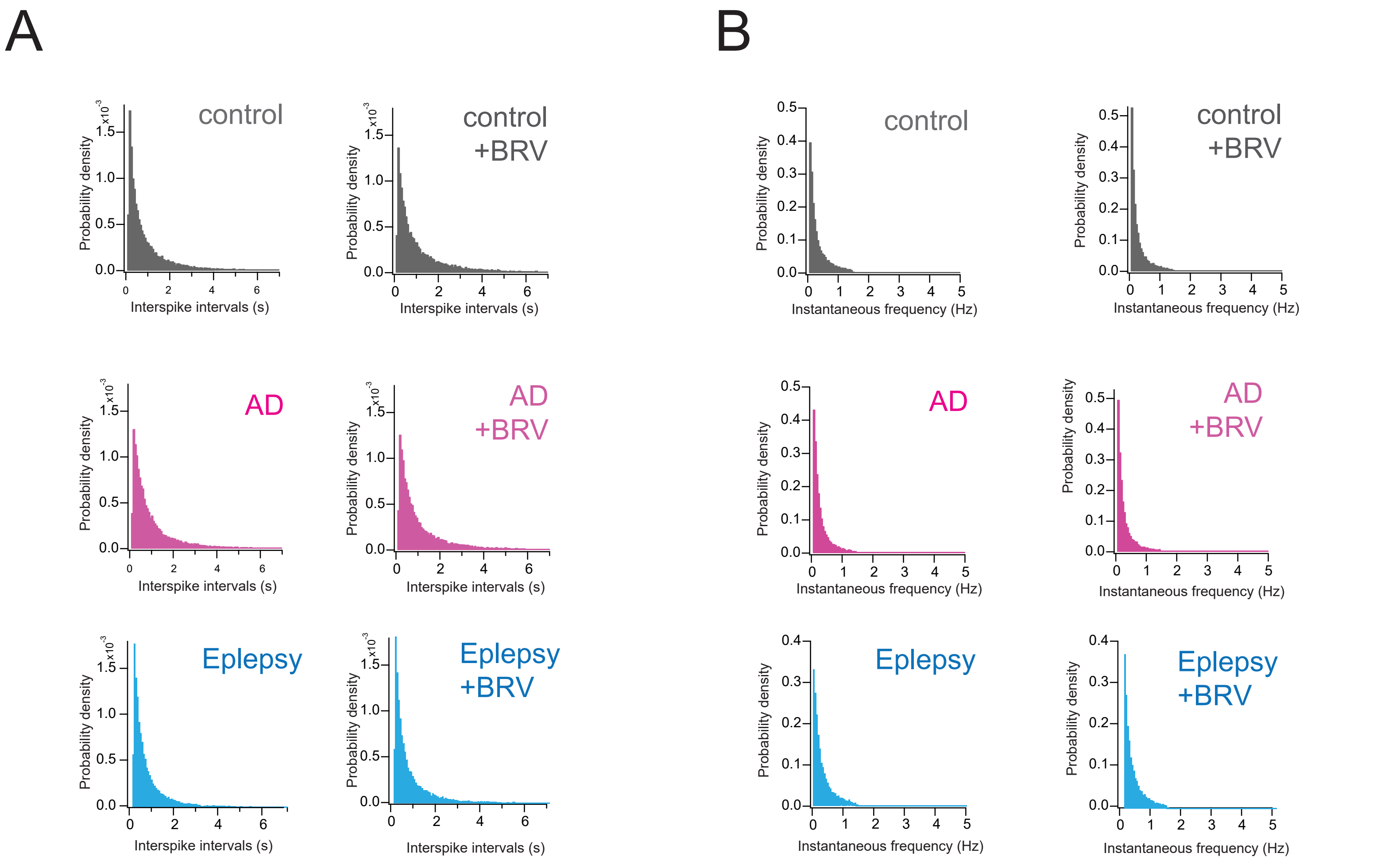
